# A chemical biology view of bioactive small molecules and a binder-based approach to connect biology to precision medicines

**DOI:** 10.1101/386904

**Authors:** Stuart L. Schreiber

## Introduction

A simplistic view of drug discovery is that it begins, most often using “model organisms”, with biological inferences of a disease that suggest the need to interfere with some activity, function or process. An enzyme should be inhibited or a pathogen should be killed. Chemical experimentation yields the desired inhibitor, and clinical investigations then test the underlying hypothesis in humans. If the stars align, an effective drug emerges.

The high cost of testing and low rate of success of this paradigm has led the drug-discovery enterprise to search for new and more effective approaches. In this essay, I explore a concept that focuses on the discovery of **compounds that bind targets** rather than inhibit a biochemical activity. I present evidence that such ‘binders’ can affect protein activity in under-appreciated ways that have therapeutic potential. This approach is well aligned with a current trend in drug discovery – to exploit insights from human biology in order to select therapeutic targets with greater confidence and to understand the deficiencies of the targets that need correction. The targets emerging from this approach most frequently lack the “simple” activities that drive much of past drug discovery. Compounds are needed that engage these targets in new and challenging ways to elicit the novel activities suggested by human biology, especially enhancing functions of proteins and conferring new (neo) functions to proteins (**Figure 1**).

**Figure 1.**
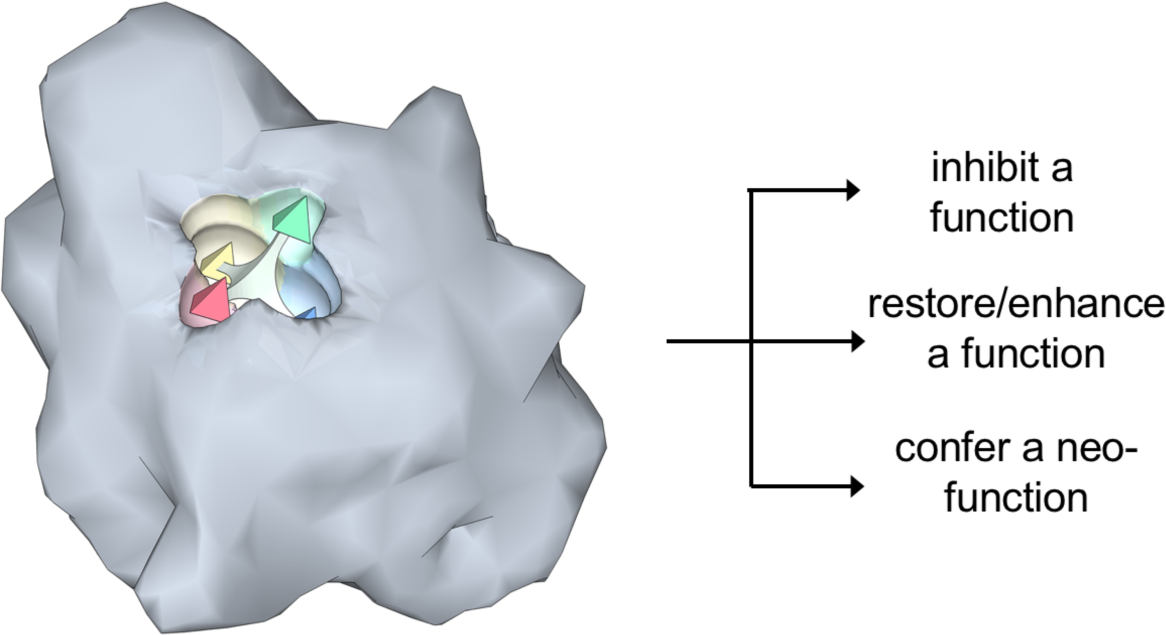
Topographically complex hot spots on proteins can be liganded with suitably shaped, often 3-D small molecules, which results in modulating functions in different ways. Binding alters the dynamic and structural features of proteins, resulting in: 1) novel interactions with other proteins and 2) changes in protein dynamics, stability, turnover rates, and tendency to be chemically modified by cellular enzymes. Each of these underappreciated effects can have therapeutic consequences. In addition to the common use of binders to inhibit function, they can also restore or enhance function, or even create a new function. Achieving a framework of chemical biology that emulates nature and evolution, where nature evolves and optimizes not so much by losing functions, but by enhancing functions and inventing new ones, promises to unlock potential not only for eliminating disease states but also for enhancing and augmenting states of health and wellness.

Assuming the challenging nature of emerging targets can be overcome, human biology-guided target selection, made possible by powerful new capabilities (low-cost, massive-scale DNA sequencing; reprogramming and editing of human cells; etc.), is expected to improve our ability to identify relevant targets for therapeutic intervention and thus overall to advance new “precision medicines”.

### A quick review of human genetics in drug discovery

Associating natural variants of human genes with health and disease can yield multiple series of alleles having low to high influence on disease. Correlating gene activity with the associated risk for or protection from a disease provides a relationship akin to a dose–response plot relating gene activity to disease.^[1]^ This dose–response is grounded in human physiology, and thus provides insight into genes encoding candidate therapeutic targets even prior to undertaking a drug-discovery effort. Linking these relationships of gene activity to other elements of human health through electronic medical records theoretically can even provide insights into the safety of hoped-for drugs (**Figure 2**).

**Figure 2.**
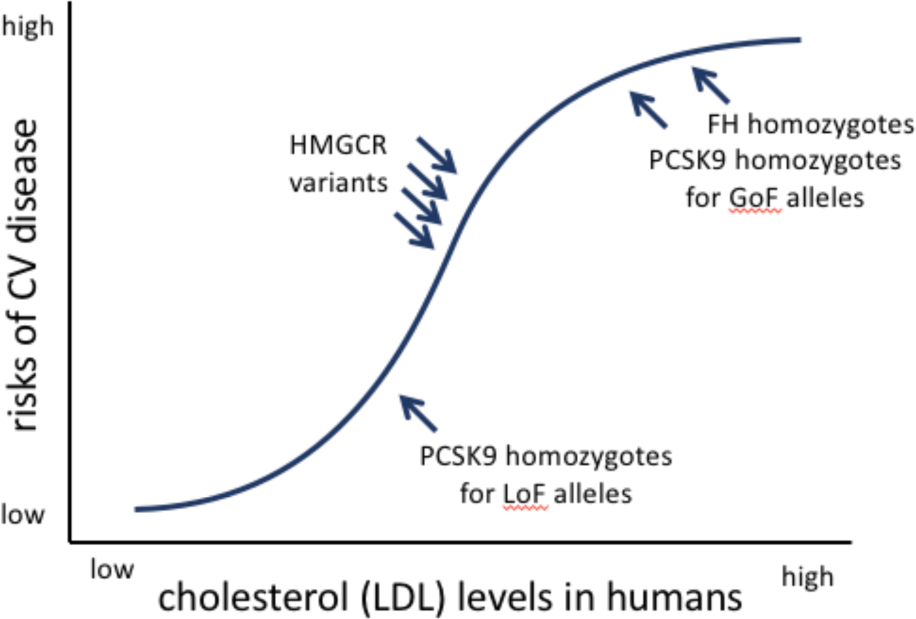
Analyses of human variants of three genes associated with risk of cardiovascular disease and altered levels of cholesterol support what we now know to be true – drugs that mimic the protective variants lower LDL levels and are heart protective. Related prospective analyses in many diseases are suggesting the activities that safe and effective drugs should confer on novel targets in the context of human physiology and prior to the start of a drug-discovery effort, but these activities are often unfamiliar to traditional efforts and may require new approaches to drug discovery. (Adapted from Reference 1.)

But gene activity is only a proxy for protein activity, and proteins are the most common targets of drugs. A hard but essential next step in human genetics-guided drug discovery is to understand the biochemical mechanisms of variant proteins encoded by disease alleles. This key but often difficult-to-obtain understanding can provide a blueprint for the precise activities that drugs should confer on their targets in order to be safe and effective. Most of this approach is still aspirational, but the few available examples, including the antibody evolocumab, a PCSK9-binding cholesterol-lowering protein therapy, are impressive and encouraging.^[2]^ PCSK9 is an extracellular protein and the therapeutic antibody evolocumab emerged from inferences derived from PCSK9 risk and protective alleles. This essay takes its inspiration in part from examples like evolocumab, but seeks to generalize the approach, especially to the more common and highly therapeutically relevant intracellular and difficult-to-drug proteins. To do so will require the discovery of small molecules that can access these targets more readily than antibodies and, as described below, modulate functions in ways that transcend simple inhibition.

A challenging step will be to discover drugs having the novel mechanisms of action suggested by biochemical investigations of gene variants. These mechanisms include **increasing the activity of a protein** or **causing it to have a new activity** (**Figure 1**). Most often, the suggested target proteins are intracellular and lack enzymatic activity – nuclear transcription factors or cytoplasmic scaffolding proteins are common examples – and thus fall into the difficult-to-drug category. For example, if we learn that a causal variant of a transcription factor involved in heart function is expressed at half the level of the wild-type allele and increases the risk of heart disease by a factor of two, we might hypothesize that either doubling the transcription factor’s stability, lifetime or activity would offer therapeutic benefit. But how do you make a drug having one of these properties? It would seem challenging – but the example of the much simpler challenge associated with evolocumab points to a potential general solution.

### Small-molecule binders in drug discovery

The chemical biology analysis of small-molecule binding to cellular targets suggests a nontraditional path to future drug discovery. This essay offers examples of these learnings from the past three decades and proposes a binder-based path toward the discovery of compounds having the novel mechanisms of action suggested by human biology. This path provides a means to bridge the currently large gap between modern biology and its therapeutic impact on patients.

The first step to this approach is to discover compounds that bind proteins, including proteins lacking enzymatic activities. An essential premise of this proposal is that **binders have hidden magic just waiting to be revealed**.

*Observations from past studies.* There is a common view of small molecules (a.k.a. chemicals; compounds; drugs) as ‘inhibitors’. Indeed, the term ‘inhibitor’ is often used as a synonym for a bioactive compound. This language suggests a limited appreciation of the myriad consequences of compounds binding proteins in cells. These consequences have important implications for the discovery of bioactive compounds. Two common consequences of binding emphasized here are:

1. Small molecules **alter the interactomes** of their targets – including the induction of novel protein associations that rewire cellular circuitry
2. Small molecules **alter the dynamic properties and cellular stabilities** of their targets – resulting in changes in the rates of post-translational modifications and concomitantly the activities of the targeted proteins, and either prolonged or shortened lifetimes

These are common activities conferred by small molecules on their targets, and they are the activities frequently suggested by insights from human genetics.

### 1. Small molecules alter the interactomes of their targets – including the induction of novel protein associations that rewire cellular circuitry

The disruption of protein–protein interactions is a well-appreciated aspect of drug discovery. However, a common but less widely appreciated outcome of binding is the association of novel proteins to the small molecule– protein complex – that is, the interactome of the liganded protein is altered. When a small molecule binds its protein target, a composite surface or dynamic/entropic feature results that can attract new protein interactions. This is analogous to a neomorph mutation inducing a new function, most often by resulting in a novel (neo) protein interaction.

Although the notion of compounds disrupting protein–protein interactions is often conceived as a goal in drug discovery (and lamented as challenging), my guess is that induced protein associations are a far more common outcome – and one that has been demonstrated to have therapeutic consequences. This phenomenon was recognized in studies of the cellular mechanisms of action of the natural products FK506, cyclosporine and rapamycin. FK506 and rapamycin bind the protein FKBP12^[3]^, which induces neo-associations with the protein phosphatase calcineurin and the protein kinase mTOR, respectively (**Figure 3**).^[4]^ Similarly, cyclosporin binds cyclophilin and induces a neo-association with calcineurin. The consequence of these novel interactions is that a subset of the activities of the phosphatase and kinase are modulated. Later, the natural product brefeldin A was found to induce a novel interaction between the guanine nucleotide exchange factor GBF1 and the guanine nucleotide-binding protein Arf1p, altering the function of GBF1.^[5]^ Analogously, the natural product abscisic acid binds the abscisic acid receptor PYR/PYL/RCAR and induces a neo-association with the protein phosphatase PP2C, again altering the function of PP2C.^[6]^ Naturally occurring phorbol esters offer a twist on neo-associations – upon binding C1 domains of protein kinase C paralogs, the kinases are induced to associate with the inner leaflet of the plasma membrane.

**Figure 3.**
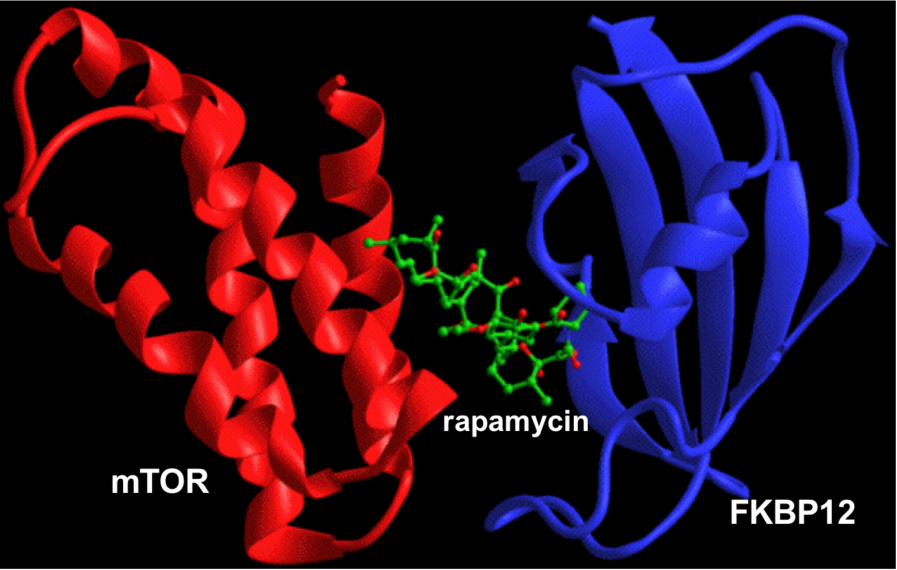
FKBP12 and mTOR do not associate with each other, and rapamycin does not bind mTOR – the mechanistic target of rapamycin. Rapamycin binds FKBP12 (blue) with extremely high affinity (200 pM) and the resulting small molecule–protein complex binds mTOR with high affinity and specificity (the FKBP12-rapamycin binding domain of mTOR (red) is shown). This illustrates how a small molecule can induce neo-protein associations.

Initially, it appeared (at least to me) as if these examples revealed the remarkable consequences of ~ a billion years of natural selection – surely these novel induced protein interactions are highly unlikely and require an eon of evolutionary tinkering. But recent chemical biology investigations have revealed that small molecule-induced protein associations are a common feature of even very simple synthetic compounds; for example, the off-the-shelf chemicals synstab A and B induce alpha and beta tubulin interactions, analogous to the complex natural products discodermolide and taxol.^[7]^ Further, the simple synthetic compounds thalidomide and lenalidomide induce (or enhance) associations of the E3 ligase complex CUL4-cereblon with the transcription factors aiolos and ikaros. In this case, the consequence is small molecule-induced degradation of these transcription factors. This outcome illustrates the interplay of the two consequences of binding emphasized in this essay; here, binding alters the interactome of the small-molecule’s target *and* thereby alters its cellular lifetime.^[8]^ The simple sulfonamides indisulam and E7820 similarly induce association of the E3 ligase CUL4-DCAF15 with the splicing factor RBM39, with consequential degradation of RBM39.^[9]^ The simple synthetic compound DNMDP induces association of the phosphodiesterase PDE3A with SLFN12, bestowing a novel function upon PDE3A that selectively kills cancer cells expressing high levels of SLFN12.^[10]^

A related consequence of binding is the selective reduction in protein–protein interactions involving scaffolding proteins; for example, the discovery of a CARD9 binder that mimics a Crohn’s Disease protective allele by removing only the TREM62 protein from the CARD9 multimeric complex.^[11]^ These and other studies suggest that bioactive small molecules commonly alter the interactomes of their protein targets; indeed, it is possible that small molecule-induced protein associations are the norm rather than the exception. This insight provides a novel proactive approach to modulating cellular functions suggested by human genetics. To exploit this insight, we’ll want to discover binders to therapeutic targets and assess; for example, using quantitative proteomics, the induced changes in target interactomes.

This last speculation is supported by the many novel gain-of-function activities of ‘chemical inducers of dimerization’, which rely on fusing dominant small molecule-sensitive dimerization domains to several hundred cellular proteins.^[12]^ These agents were inspired by the observations of natural product-induced protein associations^[13]^ and have been shown to lend small-molecule control of a wide range of biological processes, including trafficking, gene expression, protein degradation, and apoptosis, among many others. More recently, this concept was extended to enable small-molecule control of the stability of proteins of interest fused to a destabilizing domain (for example, allowing bilirubin control over target protein stability and half-life).^[14]^ A related approach enables dumbbell-shaped small-molecule degraders to induce the degradation of native proteins by exploiting the mechanisms of thalidomide and lenalidomide described above.^[15]^

Exploiting the ability of small molecules to induce new protein associations is a promising and likely general method to impart novel mechanisms of action on small-molecule probes and therapeutics. This approach is well positioned to address the challenges of drugging the targets and processes that are being illuminated by human genetics.

### 2. Small molecules alter the dynamic properties and cellular stabilities of their targets – resulting in changes in the rates of post-translational modifications and either prolonged or shortened lifetimes

In the early 1990s, my lab was studying the structure and function of SH3 domains found in numerous signaling molecules. We were able to express numerous recombinant SH3 domains alone in cells, but we noticed that they were often on the edge of folding stability.^[16]^ For example, we observed that the majority of backbone amide NHs was rapidly exchanged in D_2_O. Our project focused on the binding mode of SH3 domains, revealing their preference for proline-rich 3^10^-helical peptides, and the discovery of SH3 domain binders, which was achieved using biased combinatorial libraries.^[17]^ But we were surprised and impressed by the ability of our binders to stabilize the SH3 domains. The NH to ND exchange in D_2_O was dramatically decreased and limited to only a small number of backbone NHs.

During this period, my lab also serendipitously observed a case of small molecule-induced alteration in a post-translational modification of a targeted protein in cells. The small molecule wortmannin prevents the autophosphorylation of mTOR at Ser-2481, while the direct mTOR inhibitor rapamycin, through its intracellular FKBP12 complex, fails to do so.^[18]^ It seems to me reasonable to conjecture that binding alters protein dynamics and stability, and that this consequence of binding may alter rates of post-translational modifications of proteins or their ability to traffic or function. That the latter two are relevant to drug discovery has been established through the development of revolutionary cystic fibrosis drugs at Vertex – CFTR binders that function as correctors (binding leads to greater stability and thus efficiency of trafficking through the secretory pathway; **Figure 4**) or potentiators (binding leads to activation by altering protein dynamics).^[19]^ And since the functions of many proteins are regulated by post-translational modifications, this may be another avenue to discover compounds having novel mechanisms of action – ones not involving direct enzyme inhibition.

**Figure 4.**
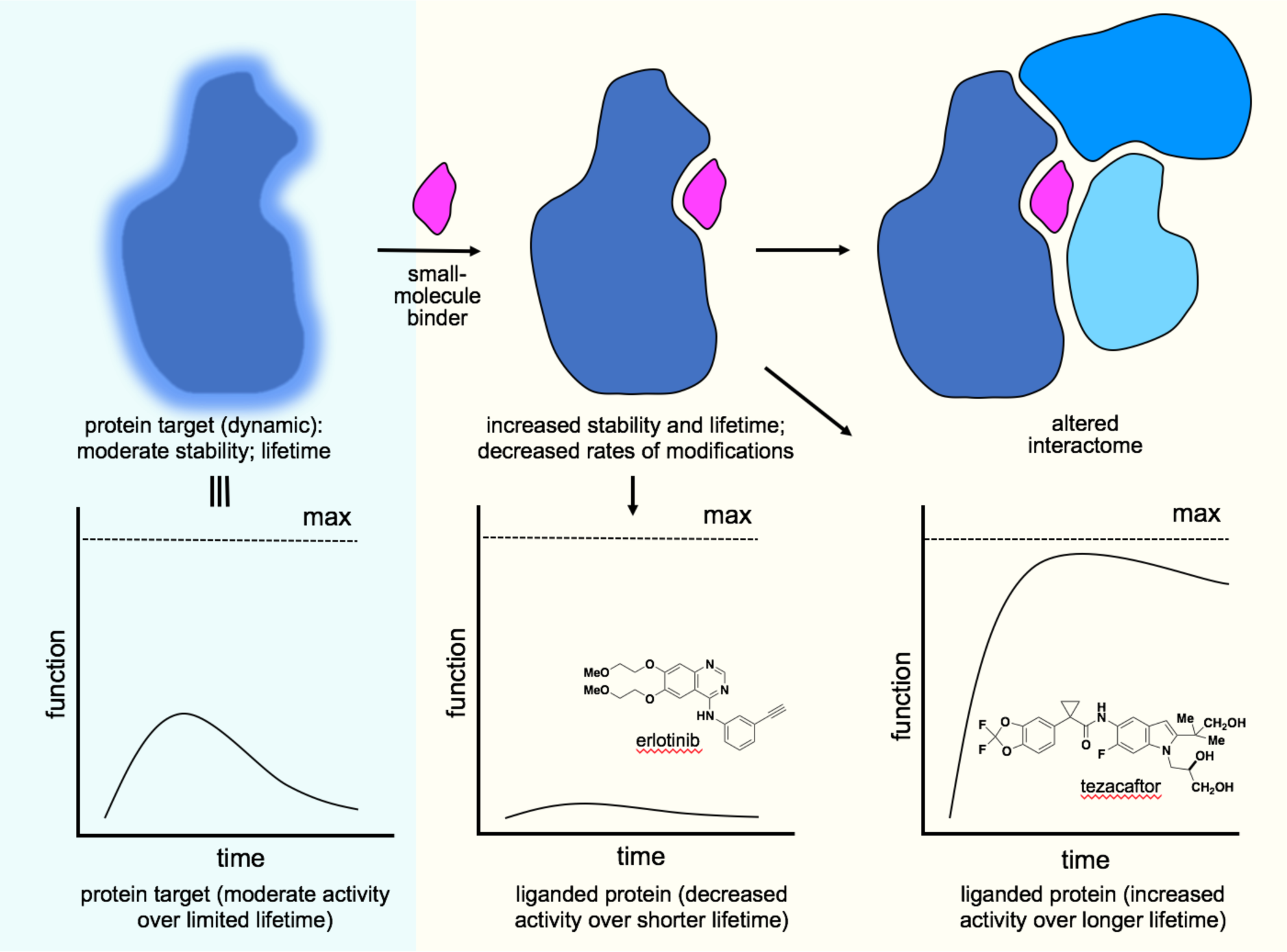
Conceptual outline of a chemical biology view of bioactive small molecules. Small-molecule binding alters the dynamic properties of proteins, often pre-organizing the protein for neo-protein interactions and/or reducing the entropic cost of binding. The ability of small molecules to alter protein interactomes; post-translational modifications; cellular turnover rates and lifetimes (e.g., erlotinib, which functions as a degrader of its target EGFR); trafficking to the functionally relevant cellular compartment (e.g., tezacaftor, which functions as a translocator of its target CFTR); among others lead to the modulation of activities relevant to drug discovery. This suggests a path to drug discovery beginning with the discovery of binders followed by the systematic determination of the cellular consequences of binding.

Since small-molecule binding to proteins alters the protein’s dynamic properties, it follows that binding should change the stability of proteins in cells and therefore their cellular lifetimes. This is what my lab observed when we first discovered SH3 domain binders – adding binders to cells increased the lifetime of the recombinantly expressed domains in cells. Evidence of the relevance of this concept to drug discovery is seen in the realization that many “kinase inhibitors”, including erlotinib and gefitinib, actually induce protein kinase degradation upon binding (EGFR in the case of erlotinib; **Figure 4**), which calls into question the MoA of even simple enzyme inhibitors (there is no kinase activity to inhibit if the kinase is absent from binding-induced degradation).^[20]^ Several binding assays, including the powerful “Drug Affinity Responsive Target Stability (DARTS) assay, rely on the principle that small-molecule binding alters protein dynamics – in the case of DARTS resulting in protease resistance.^[21]^ A longer-lived protein contributes a greater degree of its function than a shorter-lived equivalent. Indeed, there is now an abundant body of evidence that small-molecule binding leads to either increased or decreased stability of the liganded protein. This has been observed repeatedly using the extremely powerful and widely used “Cellular Thermal Shift Assay” (CETSA).^[22]^

CETSA experiments are typically performed to provide evidence of target engagement in cells. Binding often results in increased thermal cellular stability, as might be expected, but in some cases, results in decreased thermal cellular stability. A common rationale for the latter is that compounds bind less stable conformations of proteins. This empirical finding opens an avenue for finding different binders to the same protein that either extend or diminish the lifetime of activity of its target. This insight provides another novel approach to modulating cellular functions, especially ones suggested by human genetics. To exploit this insight, we’ll want to discover binders to therapeutic targets and assess induced changes in cellular stability, lifetime and turnover rates, for example (in the latter case) by using classical S^35^-methionine-incorporation pulse–chase experiments or other more modern variants, including systematic approaches that may assess changes proteome-wide.

### CONCEPT: An alternative approach to discovering human biology-guided therapeutic agents in the future: the use of binders that alter interactomes, protein modifications, cellular lifetimes, and ultimately the specific functions of proteins relevant to human health (Figure 4)

Relying on insights from human biology for selecting targets, and creating a blueprint for the activities that drugs should confer on those targets, is an appealing starting point in drug discovery. We are able today to point to many such ‘experiments of nature’. We can gain relevant, but not perfect, insights into the consequences of increased and decreased gene activity in the context of human physiology. Scanning medical records may enable insights into this kind of dose–response across many facets of human physiology, thus illuminating elements of both safety and efficacy. Although the activities suggested thus far are nontraditional, and on the surface challenging, the nontraditional view of bioactive small molecules presented here provides a two-phase path forward to the early phase of drug discovery – the identification of compounds that function by the suggested novel mechanisms of action. This approach contrasts with two of the most common means of discovering bioactive compounds: through the use of biochemical/enzymatic activity or phenotypic cell-based assays. Instead, it involves first, novel assays to discover binders, and second, cellular assays to assess the effect of binding on cellular function. These two phases of this concept for drug discovery are explored below.

### Phase 1: A Platform to discover validated small-molecule binders of proteins implicated in disease

Surprisingly, there currently exists only a limited capability for discovering small-molecule binders – either to proteins or RNAs. Fragment-based screening (FBS) is one method, but it is most often challenging to transform the weak initial binders to the requisite highly specific and potent binders, and the idiosyncratic manner of optimization makes it difficult to imagine that FBS can evolve to even a medium-throughput process.^[23]^ A major advance that promises to transform this approach enables the discovery of fragment-sized binders systematically and in intact cells using quantitative chemical proteomics. This innovation has the potential to identify binders for both specific states of proteins and protein complexes, including ones that are challenging to purify – a requirement for the traditional fragment-based approach.^[24]^

Of course, there are other promising methods for discovering binders, but I’ll focus here on another emerging one that involves synthesizing DNA bar-coded compounds – “DNA-Encoded Libraries” or DELs.^[25]^ Although this discovery technique is still in its development phase, and one could fairly say the promise outweighs the number of successes, it appears to be a method that could evolve to be robust, reasonably high throughput and, most appealingly, significantly enhanced by recent advances in synthetic organic chemistry methodology and strategy.

In traditional biochemical activity- and cell-based screens, the value of including compounds with 3-D stereochemical and structural features often found in natural products has recently been summarized (**Figure 1**).^[26]^ Compounds arising from modern asymmetric synthesis, and proven to have a novel mechanism of action, stand in contrast to the typical compounds thus far incorporated into DELs, even more so than the compounds that populate commercial vendor libraries for traditional screening. A reasonable proposal is that the value of DELs would be enhanced if a far greater range of candidate binders could be synthesized within the constraints of DEL technology, which include the need to perform synthetic transformations that are compatible with DNA and water. Nevertheless, impressive gains have recently been reported, and it seems likely that synthetic chemists will master this new challenge.^[27]^ As a lab that is coming to this field later than many others, we have already achieved several of our goals, including synthesizing DNA bar-coded compounds related to those shown in **Figure 5**.

**Figure 5.**
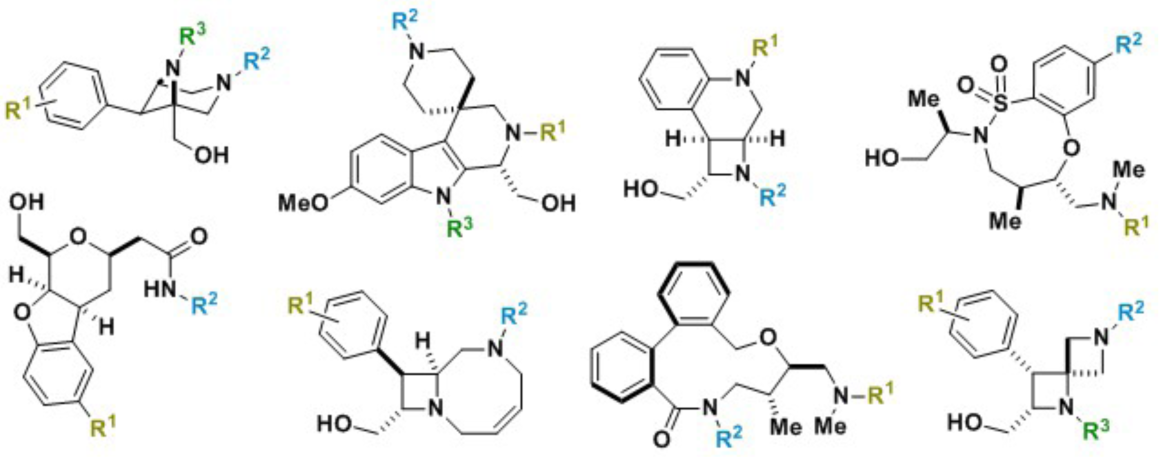
Synthetic organic chemists are innovating methods to enable modern methods of asymmetric synthesis and strategies for short syntheses of compounds having chemical features of natural products and successful probes and drugs (eight representatives shown) to be compatible with concurrent DNA-bar coding.

Whether through FBS, DELs or other strategies, the approach to drug discovery described here will require the development of a robust, high-throughput and effective means of discovering and validating binders. This Platform should include rapid and cost-efficient syntheses of putative binders. In the case of DELs, there is the added burden of resynthesizing compounds without the DNA bar-codes and testing whether they also bind. After all, the primary assay only tests binding of large (~35,000 MW!) compounds, which are overwhelmingly composed of DNA molecules (minus the ~ 400 MW small-molecule components). Time will tell whether the promise of DEL screening can override its intrinsic short-comings. In all approaches, biochemical methods will be needed to validate and quantitate the binding process to complete the first phase of the overall approach, but fortunately a plethora of effective biophysical techniques are available and likely up to this task.

### Phase 2: A Platform to determine the consequences of binding in cells and organisms, or to engineer desired features into the binders

The discovery of binders is just a first step – next is to determine whether the ‘magic of binders’ I suggested earlier can be realized. A second Platform will be needed to analyze several properties of validated binders. Staying aligned with the examples described above, it would be useful to perform interactome studies of target proteins in the free vs. liganded state. Pull-down experiments followed by quantitative proteomic analyses should provide an unbiased analysis of the ability of a given binder to induce protein associations or to alter post-translational modifications of value to specific drug-discovery efforts. Two powerful methods are available to assess the ability of binders to alter the stability of targeted proteins: 1) H/D exchange of backbone amide NHs; and 2) CETSA experiments of binders in cells, which requires simply Western blotting of the target proteins. Pulse–chase labeling experiments should be effective in assessing whether binders alter turnover rates and shorten or lengthen the half-lives of targeted proteins, and studies that correlate CETSA shifts with protein half-life might even give medicinal chemists some guidance regarding the cellular thermal shifts for which they might aim.

The pipeline above describes a plausible approach to discovering compounds that confer on their targets activities recommended by human genetics. Of course, this is simply the earliest step in an overall challenging and costly drug-discovery process; nevertheless, having the right target and modulating it in ways suggested by human biology is an important place to begin.

To illustrate, I return to my earlier theoretical example of cardiovascular disease and diminished heart function associated with diminished activity of a transcription factor. When human experiments of nature reveal the need to increase the activity of a transcription factor, and the process described above yields compounds that stabilize the transcription factor and increase its lifetime, biological experiments will of course be needed to determine whether the more stable and longer-lived transcription factor is able to correct the deficient function of a risk allele and to provide the increased function of a protective allele. When human biology-based studies reveal the need to decrease the activity of a transcription factor, analogous experiments can be performed, focusing on decreasing stability and lifetimes. These experiments are in principle informed by CETSA–stability/lifetime/function relationships that might enable predictions of the magnitude of binder-induced temperature shifts required to confer the amount of functional modulation suggested by experiments of nature. In addition, chemical inducers of dimerization including “direct degraders” like erlotinib, indisulam, E7820 and thalidomide or the new class of dumbbell-shaped degraders described above teach us that binders can possess, or can be further engineered to yield, bifunctional properties with therapeutically beneficial consequences.

## Conclusion

I started with the simple example of a promising novel mechanism of action, the cardiovascular drug evolocumab, an antibody that binds and blocks the activity of the extracellular protein PCSK9. Alleles of PCSK9 were previously identified that provided a dose–response of PCSK9 gene activity, correlating LDL levels with risk of cardiovascular disease. An antibody binder was then identified that mimicked protective alleles of PCSK9, and a powerful new therapeutic agent emerged. This illustrates a binding-based approach to drug discovery – an antibody binder mimicked the heart protective effects of a loss-of-function allele. But this example is also limited in scope – how would we apply this concept to drug targets not accessible to antibodies; to therapeutic blueprints that require the target to gain new molecular associations or to function in a more durable or more active manner; etc.?

My goal in this essay is to point readers to a possible solution to these challenges – a novel drug-discovery path that bypasses traditional biochemical activity-based screens or cell or animal-based phenotypic screens. I am imagining that this path is complementary to the two well-established ones – it may be well suited for the types of challenging non-traditional targets emphasized in this essay, whereas there will be drug-discovery efforts, for example those requiring “simple” enzyme inhibitors, that may be adequately served by the traditional approaches. The binding-based path will benefit from innovations in organic synthetic chemistry and chemical biology and provides a conceptual framework for a more general approach to drug discovery – extending to small-molecule drugs that target critical regulatory proteins that function inside of cells and that are therefore less easily modulated with protein therapeutics, such as evolocumab.

